# Root exudation facilitates water infiltration and rewetting of dry soil

**DOI:** 10.1101/2025.03.26.645447

**Authors:** Emma Gomez Peral, Andrew Mair, Iker Martin Sanchez, Gloria de las Heras Martinez, Mariya Ptashnyk, Lionel X Dupuy

## Abstract

The way plant roots facilitate water infiltration in soil may be just as important as the efficiency with which the root system in turn extracts it from the soil. Here we studied the mechanisms through which the root system facilitates water infiltration through a dry soil layer. Dye tracing experiments were conducted in model soil microcosms to characterise how root growth and exudation affects the permeability of dry layers of the model soil. Results showed that dissolved root exudates may be the primary facilitator of water infiltration, which may be linked to water surface tension. We conclude that in dry soil, root architecture and root exudation may combine to facilitate the infiltration of water and decrease the water lost by evaporation. These findings could enhance our understanding of the traits that provide drought resistance in crops.

## Introduction

Hydrologists have long acknowledged the significant role of vegetation in influencing water movement in soil, and as Clothier and Green aptly expressed, the roots are the “big movers” (Clothier and Green 1997). A primary driver is the extraction of water from the soil through root uptake and subsequent transpiration, which not only reduces soil water content but also causes water to move along gradients of the water potential. However, prior to uptake, the roots also influence water transport through modifications of the soil hydraulic properties. The presence of plant roots in soil has been linked to preferential flow of water in dye-tracing experiments. For example, Jačka et al. (2021) observed increased dye concentration around plant roots in a tracer experiment. This finding was corroborated by measurements of saturated hydraulic conductivity in soil column experiments, which showed that the soil conductivity increased due to the presence of roots (Leung et al. 2018). Various studies have also demonstrated that the hydraulic conductivity increases with the root length density (Xiao et al. 2024). In field experiments, it was also observed that most types of vegetation increase the infiltration rate of bare soil (Marshall et al. 2014; Song et al. 2017).

Changes in soil hydraulic properties can be attributed to physical modifications in the soil’s pore structure. As roots grow, they displace the surrounding soil, creating a cavity and rearranging the distribution of the surrounding soil pores (Dexter 1987). Because the size and connectivity of soil pores determine the resistance of the flow of water, these changes in soil porosity in turn affect the permeability of the soil (Schulz et al. 2019). In the study of (Anselmucci et al. 2021) roots grown in sand created an increase in porosity because of dilation. In more structured soil however, the expansion of the root cavity compacted the surrounding soil, and this can lead to a reduction in the soil porosity (Bruand et al. 1996; Helliwell et al. 2017; Koebernick et al. 2019), although this usually occurs in the bulk soils which initial porosity is low (Lucas et al. 2019). Other factors such as the shrinking or decay of the root can also create macropores (Ni et al. 2019; Duddek et al. 2022) where the flow of water can occur with least resistance.

Chemical modifications of the soil solution caused by rhizodeposition can influence water movement in soils as well. High molecular weight polymers such as mucilage increase the viscosity of the soil solution (Read et al. 1999) which in turns reduces the flow speed and increases the resistance to water movements. Mucilages behave as hydrogels because they contain polymers that are hydrophilic, crosslinked, and possess the ability to absorb and retain a significant amount of water (Naveed et al. 2019). This property is critical to maintain moisture around the root when the soil is drying (Carminati et al. 2010). Once dry however, root rhizodeposits can become water repellent and may slow down water infiltration during rewetting (Ahmed et al. 2016). Rhizodeposits also contain surfactants, and this likely facilitates water movement during infiltration and the rewetting of dry and water repellent soils (Read et al. 1999, 2003).

This study aims to determine whether physical or chemical modifications of the soil substrate have a greater impact on soil water infiltration. We carried out dye tracing experiments in model transparent soil microcosms containing dry hydrophobic layers and combined image analysis and mathematical models to quantify the respective role of root exudation and root growth on the permeability of the dry transparent soil layer.

## Material and methods

### Plant material and germination

Seeds of winter wheat (*Triticum aestum var. Filon*) were first hydrated for one hour in sterilised water and then sterilised for 15 minutes in a 1% calcium hypoclorite solution (042548.30, Thermo Scientific, Spain). After numerous washes, seeds were placed in distilled water for two more hours before transfer to agar plates in an incubator at 23 ºC. The incubation time was three days when running microcosm experiments and five days for root exudate extraction experiments.

### Extraction of root exudates

Seedlings whose radicles had reached 3-5 cm were placed on a 3D printed punched plate (5 mm diameter holes arranged on a regular grid with 2 cm spacing). Each plate hosted approximately 50 seeds and was placed on top of a 15×21×7 glass dish (Ikea, Spain) with roots dipping into 1 L of Hoagland nutrient solution (092621822, MP Biomedicals, Spain) adjusted to a pH of 7 using potassium hydroxide (KOH). Plants grew in a growth chamber (23 ºC, 60% humidity with 14/10 h day/night cycle) with the nutrient solution aerated using a pneumatic pump (Souslow, Spain) during seven days. To suppress the growth of contaminants and avoid root responses to light, the glass dish was painted in black on the outside. After six days of growth, the nutrient solution was replaced with 0.5 L of sterile distilled water. Roots grew 24 h in distilled water before three 50 ml falcon tubes (EP0030122232, Sigma-Aldrich, Spain) were used to collect 120 mL of solution from each glass dish. The root exudate solutions were then frozen at −80 ºC. To adjust the concentration of root exudates, falcon tubes containing root exudate solutions were freeze-dried and 2 mL of distilled water were subsequently added before pooling all the root exudates from one glass dish into a 15 mL falcon tube (EP0030122194, Sigma-Aldrich, Spain). The root exudate solution obtained was freeze-dried a second time for weighing, and distilled water was added to adjust the concentration of the root exudate solution to 1 mg mL^-1^. Root exudates were then stored frozen at −20 ºC. We ran 4 experiments with a total of 8 plates.

### Transparent soil

Transparent soil was prepared as described earlier (Downie et al. 2012). Briefly Nafion pellets (NR50 1100, Ion Power Inc, USA) were fractured to a texture similar to sand (0.25 to 1.25 mm) using a freezer mill (6850 Freezer/Mill, SPEX CertiPrep, UK) and a series of sieves. The pH of the substrate was then adjusted by washing with Hoagland nutrient solution. With a density for Nafion of approximately 1.9 g cm^-3^ and a porosity of 0.3, the estimated substrate density is 1.58 g cm^-3^.

### Microcosm preparation

Microcosm chambers were assembled following the protocol described by Liu et al. (2025) from microscope glass slides (76 × 26 × 1 mm^3^, VWR, Spain). Glass slides were bonded by 4 mm polydimethylsiloxane spacer (PDMS, SYLGARD 184, Sigma-Aldrich, Spain) following surface treatment by oxygen plasma for 15 s at 100 W (HPT-100, Henniker Plasma, UK). After surface treatment, the glass and PDMS surfaces were brought together, and pressure was applied to ensure a strong bond between all surfaces in contact. Following the assembly of the different part, the space available for placing plants and soil is 68 × 18 × 4 mm^3^. The microcosm was then filled with transparent soil layers with different water content (Figure 1A). The bottom 1.5 cm layer (approximate dry weight of 1.7 g) of the microcosm chamber was filled with transparent soil fully saturated with water. Then a 0.8 cm layer of dry transparent soil was added (approximate dry weight of 0.9 g). The dry transparent soil was obtained by heating in an oven at 105 ºC for 24 hours. The top layer was 1 cm thick (approximate dry weight of 1.1 g) of transparent soil fully saturated with water.

**Figure 1:**
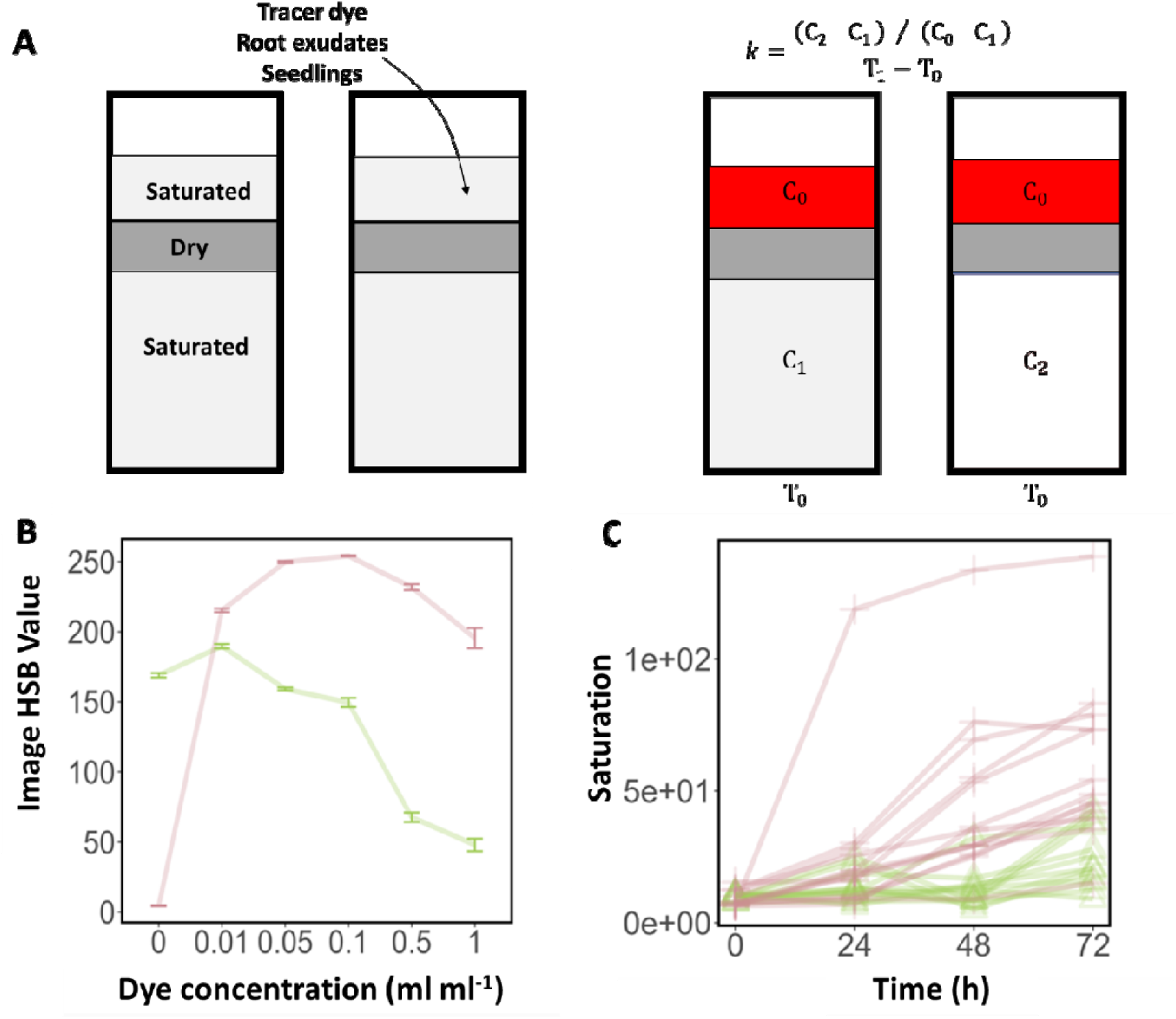
Model system for quantifying the impact of root exudates on water transport (A) Diagram of the microcosm system. At the start of the experiment, the microcosm is comprised of three layers of transparent soil (left). The top layer of transparent soil is saturated with water to allow fast diffusion of the dye. The middle layer is oven dried to produce a hydrophobic barrier. The bottom layer is saturated with water for precise determination of the quantity of dye moving through the dry layer. During the experiment (right) the dye was introduced at T0 at the top of the microcosm chamber and diffused rapidly through the top layer of transparent soil. At time T1 the tracer dye became visible, and a permeability coefficient could be calculated. (B) The dye concentration in the liquid can be quantified from the image saturation value (red) at low dye concentration, and with the image brightness value (green) at high concentration. (C) Example of time lapse data of the saturation value captured in the bottom transparent soil layer.

### Dye tracing experiment

First, we studied how the presence of plant roots affected the permeability of the dry transparent soil layer. Control samples consisted of microcosm chambers prepared as described in the previous section but without seedlings transferred into them. Wheat seedlings were transferred when the root length was 0.5 to 0.8 mm in length. All microcosm chambers were closed with parafilm tape (HS234526B, Sigma-Aldrich, Spain) and placed in a growth chamber at 23 ºC, 60% humidity with 14/10 h day/night cycle. Distilled water was added to the samples using a syringe to maintain the top layer of transparent soil saturated. The tracer dye was added 8 days after the transfer of the root seedlings. Most plants had at least one root in the bottom of the microcosm chamber at that time. A food dye (red food colorant, Vahiné, Spain) was diluted to 0.1 mL mL^-1^ to obtain the tracer dye solution. 0.2 mL of that solution was then added on the top of the microcosm chamber. Half an hour later, the first image was taken (t=0 hour). The following images were taken 24, 48 and 72 hours after the first image. The experiment was repeated 3 times with a total of 15 samples with roots and 13 controls samples without roots. Two root samples were discarded because of leakage of the microcosm chamber.

In the second experiment, we quantified the effect of root exudates on the infiltration of water. In this case the tracer dye solution was diluted with the root exudate solution so that resulting root exudate concentration was approximately 0.9 mg mL. Then 0.2 mL of that solution was then added at the top of the microcosm (concentration of root exudate approximately 0.16 mg g^-1^). All microcosm chambers were sealed with parafilm tape and placed in the growth chamber under the previously described conditions. The first images were obtained half an hour after inoculation (t=0 hour). The following images were obtained 72, 96 and 120 hours after the first image. The experiment was repeated 3 times with a total of 19 samples containing root exudates and 16 control samples containing only water. One control sample was discarded due to a leak.

In a third experiment, we studied the effect of the presence of root exudates in the dry transparent soil layer. To prepare dry transparent soil containing root exudates, 5 g of wet transparent soil was mixed 1 mL of the root exudate solution. The wet transparent soil was then left to dry at 105 ºC for 48 hours in an oven. Therefore, the resulting root exudate concentration in the dry transparent soil was similar to that of the top layer, 0.633 mg g^-1^. Four types of samples were prepared. Namely, samples that did not contain root exudates (NN), samples that contained root exudates in the middle layer (NE), samples that contained root exudates in the top layer EN), and samples that contained root exudates in both layers (EE). All microcosm chambers were sealed with parafilm tape and placed in the growth chamber. The first image was acquired half an hour after introduction of the tracer dye (t=0 hour). The following images were taken 72, 96 and 120 hours after the first image. The experiment was repeated 3 times with a total of 8 samples for each combination.

Finally, in the fourth experiment, we studied the effect of the rearrangement of transparent soil particles induced by penetration of the dry middle layer. A hypodermic needle (12684536, Fisher Scientific, Spain) was pushed approximately 2.5 cm into the transparent soil. The needle was 0.8 mm in diameter which is similar to the diameter of wheat roots (Hallam et al. 2021). Control samples were not penetrated by the hypodermic needle. 0.2 ml of the tracer dye solution was added at the top of the microcosm chamber. All microcosm chambers were sealed with parafilm tape and placed in the growth chamber. Half an hour after inoculation of the tracer dye, the first image was taken (t=0 hour). The following images were taken 24, 48 and 120 hours after the first image. The experiment was repeated 3 times with a total of 15 samples and 15 controls. One sample was discarded due to a leak.

### Quantitative image analysis

Images of microcosm chambers were acquired using a Canon EOS 2000D camera (Canon, Spain) with an exposure time of 1/1000 s, ISO-400 speed, and focal length was 39 mm and a f-stop of f/5.6. Colour images were stored uncompressed in colour format (sRGB) at resolution of 6000 × 4000 pixels. To quantify the tracer dye concentration from image data, calibration samples with dye concentration of 0, 0.01, 0.05, 0.1, 0.5 and 1 mL mL^-1^ were prepared in triplicate. Images were acquired for each sample and relationships were established between the dye concentration and the image brightness, and between dye concentration and the image saturation value (Figure 1B). To quantify dye concentration in the microcosm chambers containing transparent soil layers, regions of the image corresponding to the top, middle and bottom layers of transparent soil were cropped from the raw image data. Images were transformed to hue, saturation and brightness (HSB) format and the saturation value was used to determine the dye concentration (Figure 1C). All image processing tasks were performed using the ImageJ software (Schmid et al. 2010).

The permeability of the dry transparent soil layer *k* [s^-1^] was calculated using the following formula

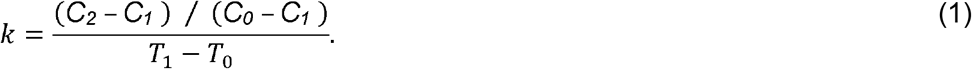

Here, *C*_0_ and *C*_1_ are the dye concentrations at time *T*_0_ in the top and bottom layer respectively. *C*_2_ is the dye concentration in the bottom layer at time *T*_1_.

### Statistical analysis

Statistical models were constructed to analyse the effects of plant roots, root exudates or to analyse the effect of the needle on both the permeability *k* and the tracer dye concentration in the dry transparent soil layer. We used generalized linear models to correlate the permeability of the dry transparent soil layer with both time and treatment effects. We tried different variance functions and found that heterogeneity was best corrected using variance covarying with treatments rather than with time. Model selection was based on the Akaike information criterion (AIC). Statistical analyses were performed with R (R Core Team 2021) using the glm package.

### Model of water infiltration and rewetting

Water transport in the transparent soil microcosms was represented using the model of Mair (unpublished). The model solves Richards equations (Richards, 1931), on a spatial domain which here is in 1D,*x* ∈[ −3.3,0], *t* ∈ (0,*T*]. Final time was T=5 days (120 hours).

To match the experimental conditions, the pressure head was adjusted to −0.01 kPa −0.1 cm at the top −100 kPa in the middle layer and −0.01 kPa at the bottom. The water content and hydraulic conductivity functions were expressed from the soil pressure head using the formulations of (van Genuchten 1980) and (Mualem 1976):

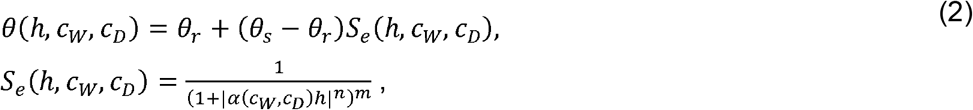

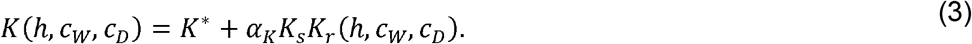

Both the Nafion particles and root exudates are known to have different behaviour during wetting and drying (hysteresis in both the hydraulic conductivity and water retention functions), and this was modelled represented using the model of Kool and Parker (1987) for water content θ (cm^3^cm^-3^)and the model of Vogel and Zhang (1996) for the hydraulic conductivity *K* (cm d^−1^). Hence, we focused primarily on the influence of the concentration of root exudates suspended in the solution *cw* (mg mL^−1^) and the concentration of root exudates dried and attached to the surface of the particles c_*D*_ (mg g^−1^). These parameters affect the inverse air entry pressure head α (cm^−1^), as described in (Karagunduz et al. 2001), through the surface tension of water *γ* (mN m^−1^) and water contact angle *ω* (rad)

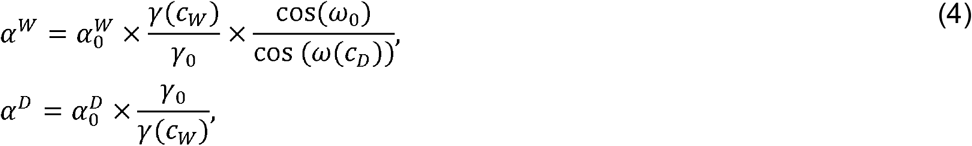

Here *α*^*W*^ and *α*^*D*^ are the inverse air entry pressure head for wetting and drying respectively and 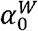 and 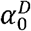 are the default inverse air entry pressures if no exudates are present in either form. The effect of root exudates is also incorporated into the saturated hydraulic conductivity as

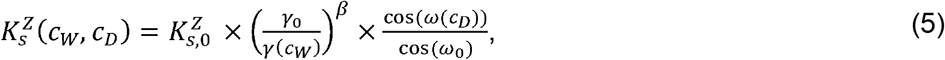

where *Z* = *W*,*D* for wetting or drying. Here 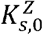 are the default saturated hydraulic conductivities during wetting and drying if no exudates are present, and *β* determines the extent to which an exudate-induced reduction in surface tension increases hydraulic conductivity. The terms *K*\* and *γ*_*K*_ in (3) are included to effectively incorporate hysteresis. The expressions for surface tension and contact angle as functions of the concentrations of suspended and dried root exudates are fitted to the data of Read et al. (2003) and Zickenrott et al., (2016) respectively,

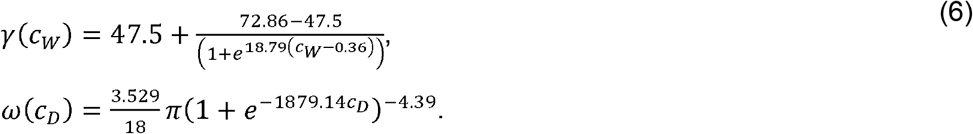

The dynamics of the root exudates in solution and the dried root exudates are given by the following differential equations

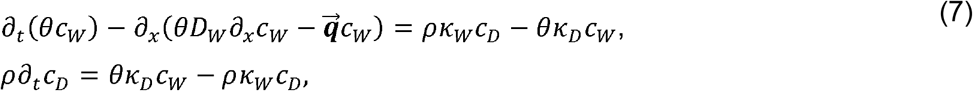

where *D*_*w*_ (cm^2^ d^−1^) is the coefficient of diffusion of root exudates through soil water, 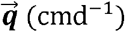 is the advective flux of soil water and *ρ* (mg cm^−3^) is the bulk density of Nafion. The terms *κ*_*w*_ and *κ*_*D*_ (d^−1^) are the rates at which dried root exudate joins the soil water solution and root exudate in solution dries to the pore surface respectively. Finally, the movement of dye within the chambers is modelled using an additional differential equation

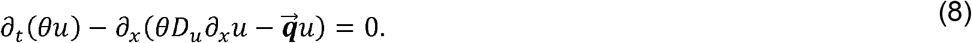

Here *u* (mL mL^−1^) is the concentration of dye in solution and *D*_*u*_ (cm^2^d^−1^) is the diffusion coefficient of the dye in solution. The dye concentration at the start of the experiment is 0.04 mL mL^−1^ in the upper layer where root exudates have been introduced, 0 otherwise.

A finite element scheme, with an implicit Euler discretisation in time, was used to approximate solutions to the coupled system of equations.

### Model parameterisation

Model parameters were taken from previously published literature assuming that coarse sand would be the best approximation of the transparent soil.

However, the reference inverse air-entry pressure heads 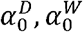 and saturated hydraulic conductivities 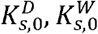 were altered considerably from the values cited in the literature in order to reflect the hydrophobicity of dry Nafion that was observed in the control infiltration experiments. The full list of parameters can be found in Table 1.

**Table 1:** Model parameter values dervied from the scientific litterature.

| Parameter | Value | Source |
| --- | --- | --- |
| $\theta_{r,0}$ | $1 \times 10^{-10} \text{ cm}^3 \text{cm}^{-3}$ | $N \setminus A$ |
| $\theta_{s,0}$ | $0.404 \text{ cm}^3 \text{cm}^{-3}$ | (Benson et al. 2014) |
| $\alpha_0^D$ | $0.049 \text{ cm}^{-1}$ | (Benson et al. 2014) |
| $\alpha_0^W$ | 0.164 cm<sup>-1</sup> | (Benson et al. 2014) |
| $n$ | 3.16 | (Benson et al. 2014) |
| $m$ | 0.684 | (Benson et al. 2014) |
| $K_{s,0}^D$ | $8.196 \times 10^{-4}$ cm d<sup>-1</sup> | (Carsel and Parrish 1988) |
| $K_{s,0}^W$ | $4.917 \times 10^{-4}$ cm d<sup>-1</sup> | (Carsel and Parrish 1988) |
| $D_W$ | 0.65 cm<sup>2</sup>d<sup>-1</sup> | (Scott et al. 1995), average taken |
| $\rho$ | 1.9 g cm<sup>-3</sup> | (Oberbroeckling et al. 2002) |
| $D_u$ | 0.5 cm<sup>2</sup>d<sup>-1</sup> | (Syms 2017) |

Two types of critical parameters could not be determined from the literature. First, the coefficient linking surface tension to hydraulic conductivity *β* is both very specific to the model system and could not be found in the literature. To calibrate the value of *β*, data was used from the second experiment and compared observations to simulated dye concentration between depths −3.3 cm and −1.8cm at times *t* = 3,4,5 d. sing Bayesian optimisation (Brochu et al. 2010) we obtained the values *β* that most accurately matched experimental results. The rates at which root exudates dissolve or reverse to a dry state are also very specific to our model system and were obtained in a similar way. Here we used the observation from the third experiment and Bayesian optimisation to determine the values of *κ*_*w*_ and *κ*_*D*_ that best predicted dye concentrations in the lower third of the domain.

## Results

### Plant roots increase the infiltration of water through dry transparent soil in a microcosm system

Following introduction into the microcosm, the dye was observed spreading non homogeneously but rapidly within the top transparent soil layer (Figure 2A), but there was no evidence of the dye being absorbed by the particles. The dry transparent soil layer blocked further spreading of the tracer dye into the lower layers of transparent soil. There was no obvious effect of the roots on the transport of the dye at this stage of the experiment. The spatial distribution of the tracer dye was analysed over a 72-hour period following the start of the experiment (Figure 2B). In the control samples, the wetting front advanced downward by 1 to 2 mm during the first 48 hours. We did not observe further progress of the wetting front during the rest of the experiment, even in the case where the tracer dye was seen in the bottom transparent soil layer (Figure 2B). 24 hours after the introduction of the tracer dye, no coloration was observed in the bottom transparent soil layer of the control samples. Only 5 samples out of 13 showed some coloration at the end of the experiment. In the samples containing roots, we observed a greater movement of the wetting front. 48 hours after the experiment began, the wetting front had advanced downward by 3 to 4 mm. When the root was visible, we observed the dye both attached to the root and in the surrounding transparent soil (Figure 2B). The tracer dye began to appear in the lower transparent soil layer 24 hours after the introduction of the tracer dye. The coloration increased in intensity over time until the end of the experiments. All samples showed some coloration at the end of the experiment.

**Figure 2:**
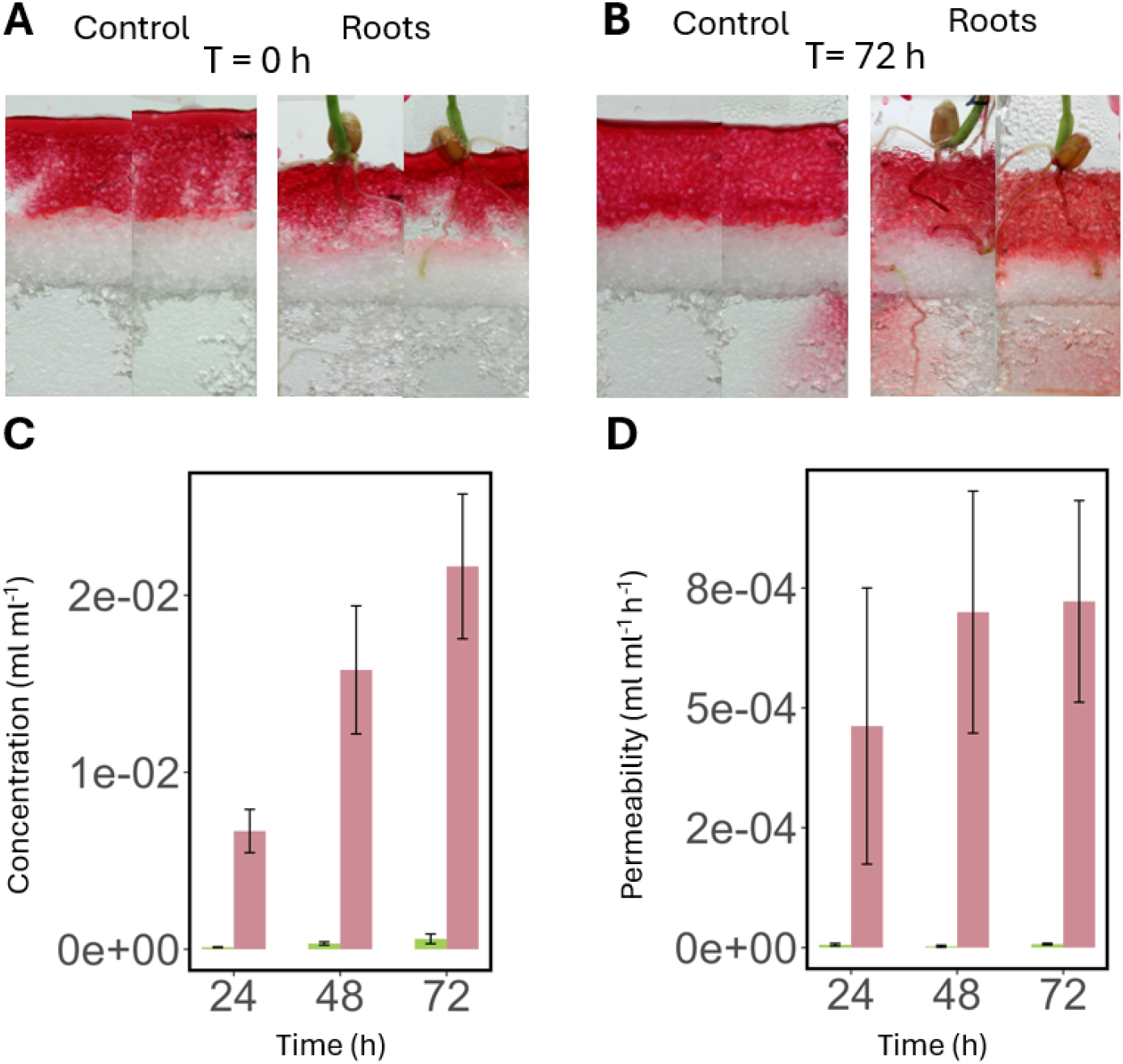
Effect of the presence of winter wheat (*Triticum aestum*) roots on the infiltration of a tracer dye through a dry transparent soil layer. **(A)** Microcosm samples just after introduction of the tracer dye into the top transparent soil layer. Sample without root (Control, left) and sample with root (right). **(B)** Microcosm samples 72 hours after introduction of the tracer dye. **(C)** Concentration of the tracer dye in the dry transparent soil layer measured at different time points. **(D)** Permeability of the dry transparent soil layer measured at different time points. C&D, Green control samples and pink samples with roots.

The quantitative analysis of the image data confirmed visual observations. The dye concentration in the dry transparent soil layer increased with time (Figure 2C). In the case of the control samples, the dye concentration increased up to a mean value of 3.5 x10^-4^ mL mL^-1^. In the case of the samples containing plants, an increase in dye concentration was observed during the entire experiment with a final dye concentration recorded at 2.1×10^-2^ mL mL^-1^. Quantitatively, the estimated dye concentration in the dry transparent soil layer increased when a root was present, but not in control treatments (Figure 2C). We did not measure a significant change over time in the permeability of the dry transparent soil layer in the control samples (Figure 2D). Statistical analysis of the data using a linear model showed that in the absence of the root, the permeability of the dry transparent soil layer was not significantly different from 0 during the entire experiment (p=0.23 and p=0.69 for slope and intercept). However, the presence of the root had a significant effect on the rate of increase in the permeability of the dry transparent soil layer (p<0.001).

### Root exudates increase water infiltration through the dry transparent soil layer

Following introduction into the microcosm, the dye was observed spreading non homogeneously but rapidly within the top transparent soil layer (Figure 3A), but there was no evidence of the dye being absorbed by the transparent soil particles. The dry transparent soil layer blocked further spreading of the tracer dye into the lower layers of transparent soil. There was no obvious effect of the presence of root exudates on the transport of the dye at this stage of the experiment. The spatial distribution of the tracer dye was analysed over a 120-hour period following the start of the experiment (Figure 3A&B). In control samples, the wetting front did not advance during the first 72 hours but had advanced by 1-2 mm by the end of the experiment (Figure 3A&B). 96 hours after the introduction of the tracer dye, red coloration began visible in the bottom transparent soil layer in all samples. The coloration increased in intensity with time until the end of the experiments. In the samples with root exudates, the wetting front advanced irregularly but continuously into the dry layer throughout the experiment. At the end of the experiment, the dry transparent soil layer was completely wet (Figure 3A&B).

**Figure 3:**
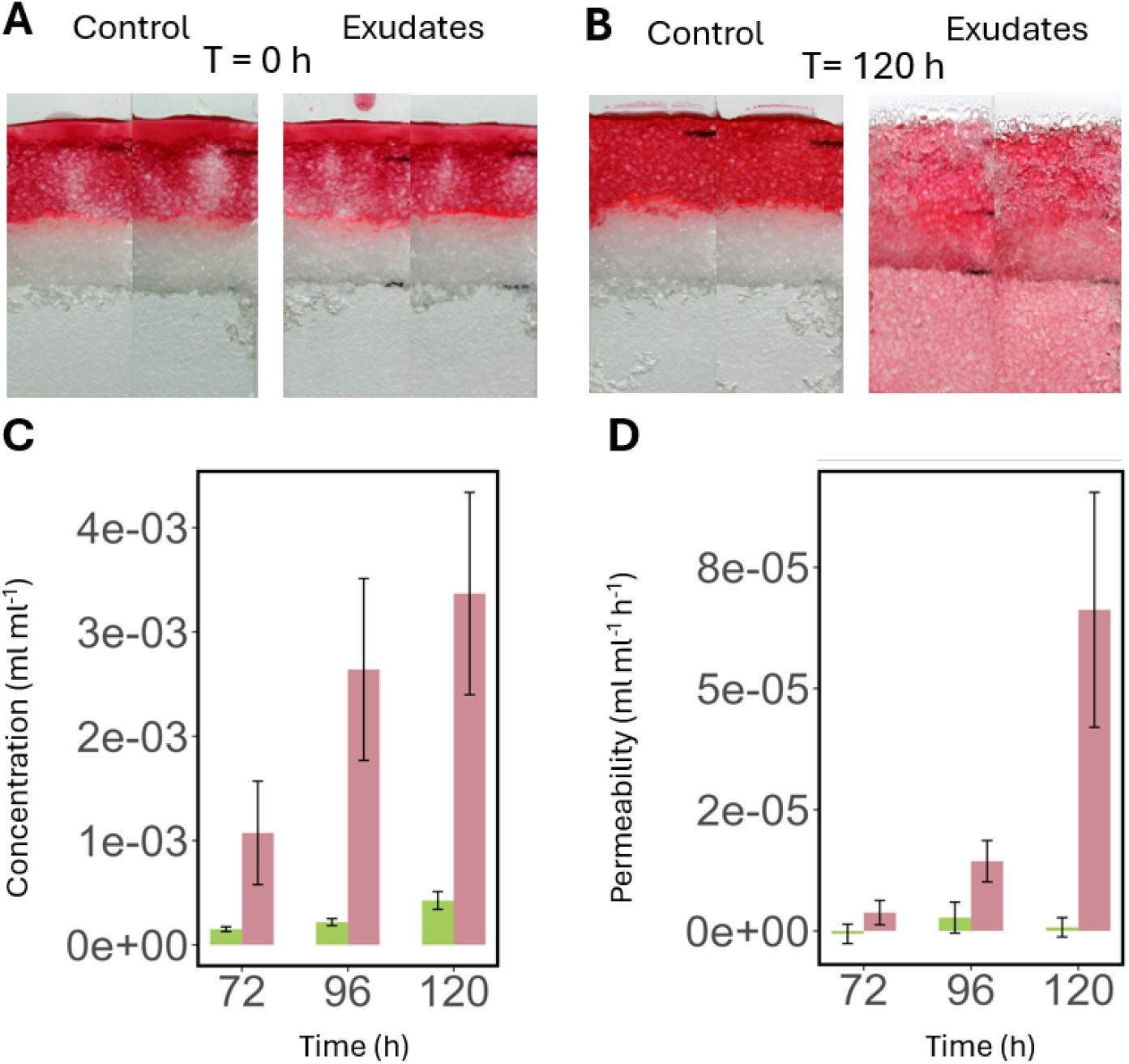
Effect of the presence or absence of root exudates from winter wheat (*Triticum aestum*) on the infiltration of a tracer dye through a dry transparent soil layer. (A) Microcosm samples just after introduction of the tracer dye into the top transparent soil layer. Sample without root exudates (left) and sample with root exudates (right). (B) Microcosm samples 120 hours after introduction the tracer dye. (C) Concentration of the tracer dye in the dry soil layer measured at different time points. (D) Permeability of the dry transparent soil layer measured at different time points. C&D green control samples and pink samples with exudates.

Quantitatively, the mean dye concentration in the dry transparent soil layer increased with time (Figure 3C) for both control samples and samples containing root exudates. When samples contained root exudates, the increase in dye concentration in the bottom layer was sustained during the experiment. We did not measure a significant change with time in the permeability of the dry transparent soil layer in the control samples (Figure 3D). For samples with root exudates, permeability was maximal 120 hours after introduction of the tracer dye. Statistical analysis of the data using a linear model showed that in the absence of root exudates, the permeability of the dry transparent soil layer was not significantly different from 0 during the entire experiment (p=0.54 and p=0.45 for slope and intercept). However, the presence of root exudates had a significant effect on the rate of increase in the permeability of the dry transparent soil layer (p<0.001).

### The presence of root exudates in the dry layers has limited effect on water infiltration and rewetting of the dry transparent soil layer

Following introduction into the microcosm, the dye spread following a similar pattern as reported in the previous experiments (Figure 4A). There was no obvious effect of the presence of root exudates (in the top or dry layer) on the transport of the dye at this stage of the experiment. The spatial distribution of the tracer dye was analysed over a 120-hour period following the start of the experiment (Figure 4A&B). In samples, that did not contain root exudates, the wetting front did not advance during the first 72 hours but had advanced by 1 to 1.5 mm both in the NN treatment and in the NE treatments (Figure 3A&B). 96 hours after the introduction of the tracer dye, red coloration began visible in the bottom transparent soil layer in all samples. The coloration increased in intensity with time until the end of the experiments. No colouration was observed at the bottom of the microcosm chambers at any stage of the experiment. There were no strong visual differences between samples with and without root exudates in the dry transparent soil layer. Samples containing root exudates in the tracer dye (EN and EE) exhibited qualitatively similar behaviour to samples containing root exudates in the tracer dye in experiment 2.

**Figure 4.**
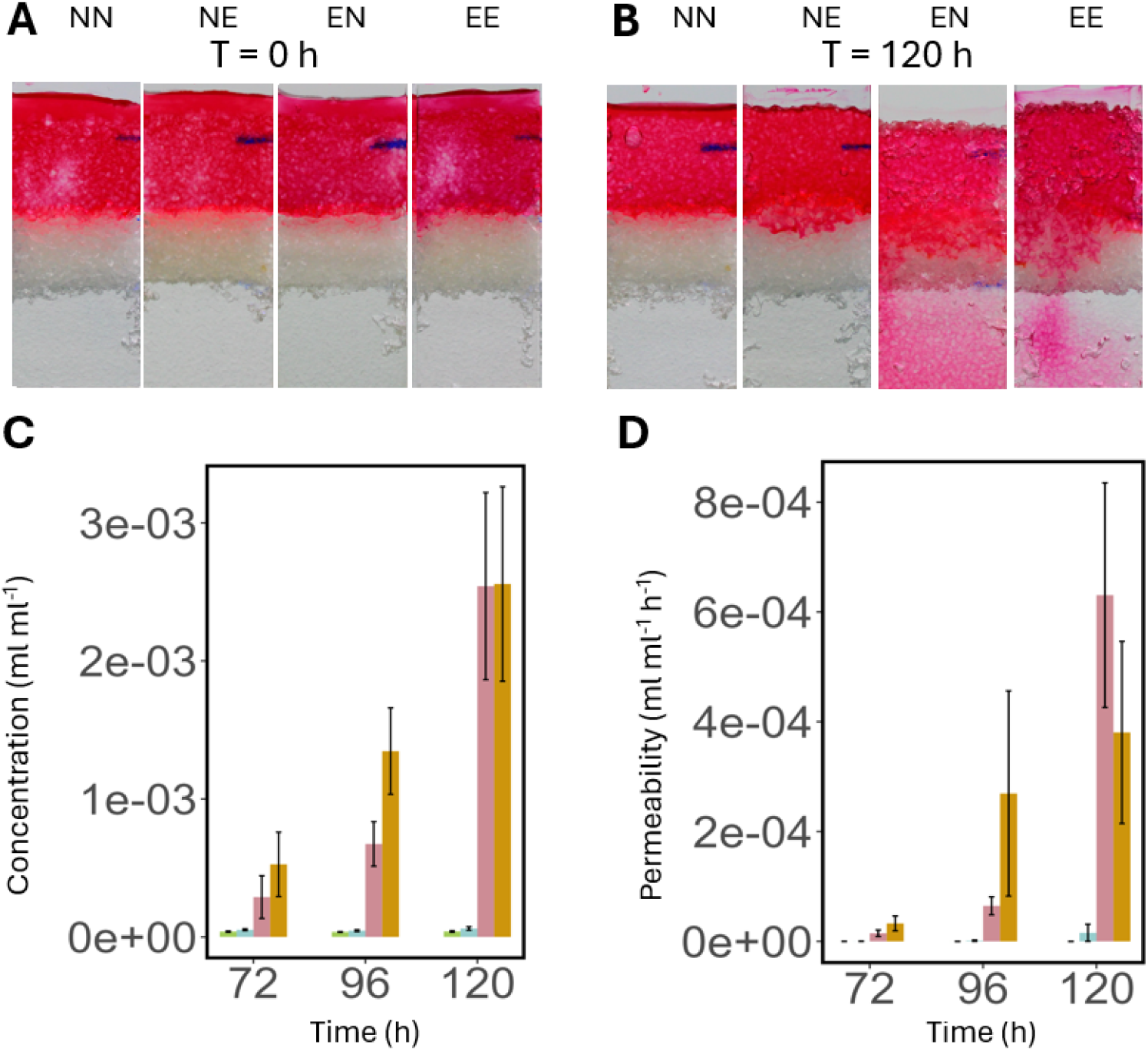
Effect of location of root exudates of winter wheat (*Triticum aestum*) on the infiltration of a tracer dye through a dry transparent soil layer. (A) Microcosm samples just after introduction of the tracer dye into the top transparent soil layer. (B) Microcosm samples 120 hours after the start of the experiment. (C) Concentration of the tracer dye in the dry transparent soil layer measured at different time points. (D) Permeability of the dry transparent soil layer measured at different time points. In C&D green NN, blue NE pink EN and yellow EE.

Quantitative analysis of the image data confirmed the visual observations. The dye concentration in the dry transparent soil layer increased with time (Figure 4C). For samples containing no exudates in the tracer dye, the increase was gradual, but for samples containing root exudates in the tracer dye, a sharp increase in dye concentration was observed within the first 72 hours and this was followed by a more moderate increase in dye concentration. We did not measure a significant change over time in the permeability of the dry transparent soil layer in samples containing water in the tracer dye (Figure 4D). In the case of samples with root exudates in the top layer, permeability steadily increased during the experiment. Statistical analysis of the data using a linear model showed that in the absence of root exudates, the permeability of the dry transparent soil layer was not significantly different from 0 during the entire experiment but the presence of root exudates in the top transparent soil layer had both a significant effect on the slope and intercept of the relationship (p<0.001 and p=0.01 respectively). The presence of root exudates in the middle layer did not have a significant effect on the permeability of the dry transparent soil layer.

### Physical modifications due to penetration of the dry transparent soil layer do not affect the infiltration and rewetting

Following introduction into the microcosm, the dye spread following a similar pattern than experiments where the tracer dye did not contain root exudates (Figure 5A&B). There was no obvious effect of the presence of the needle on the transport of the dye at this stage of the experiment. The spatial distribution of the tracer dye was analysed over a 120-hour period following the start of the experiment. No difference was observed between treatments during the entire experiment. No dye was observed at the bottom transparent soil layer, nor in the dry transparent soil layer. At the end of the experiment, we observed that the wetting front had advanced about 1 mm downward for all treatments.

**Figure 5.**
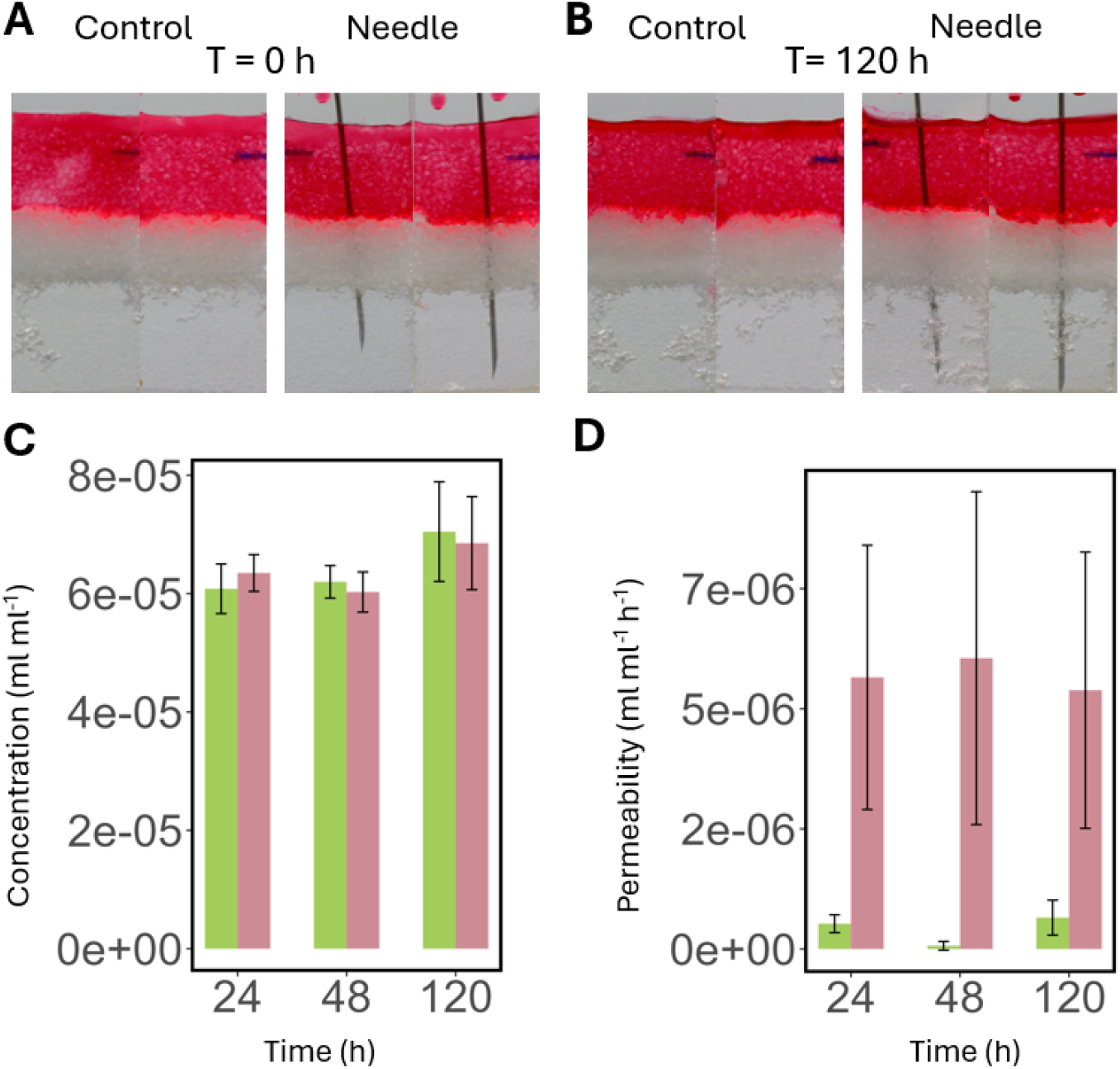
Effect of the presence or absence of a needle on the infiltration of a tracer dye through a dry transparent soil layer. (A) Microcosm samples just after introduction of the tracer dye into the top transparent soil layer. Sample without needle (Control, left) and sample with needle (right). (B) Microcosm samples 120 hours after introduction of the tracer dye. (C) Concentration of the tracer dye in the dry transparent soil layer measured at different time points. (D) Permeability of the dry transparent soil layer measured at different time points. C&D, green control samples, pink samples with a needle.

Quantitative analysis of the image data confirmed the visual observations. The dye concentration in the dry transparent soil layer did not increase with time in both conditions (Figure 5C). Calculation of the permeabilities of the dry transparent soil layer revealed that that when the needle was present, there was much higher variability and mean value for the permeability (Figure 5D), which is likely due to the needle having an effect on the saturation of the image. Statistical analyses however showed that the presence of a needle did not have an effect on the calculated permeability (P = 0.0073).

### The hydraulic conductivity is critically sensitive to surface tension and root exudate concentration

Model calibration completed successfully with optimal parameters values for 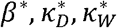 determined from experimental data. Following calibration, the predicted dye concentration was plotted against experimental data (Figure 6). Results showed good agreements between the model and experimental observations. Discrepancies were observed in the control treatment where no exudates were introduced (NN). Experiments showed an abrupt stop in the permeability of the dry transparent soil layer, which the model could not represent. The estimated model parameters gave also interesting insights into the effects of root exudates on water transport in transparent soil. The exponent of the power law relating surface tension to hydraulic conductivity was *β* = 8.8, indicating that the concentration of root exudate in solution had a strong effect on the hydraulic conductivity of transparent soil. The parameter for the solubilisation or sorption of root exudates 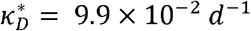 and 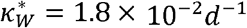 also provided useful insight. They showed that although the rate of solubilisation may not be limiting, because permeability is very small at the start of the experiments, the fraction of the transparent soil exposed to water limits the capacity of the dry root exudates to meaningfully increase the infiltration of water.

**Figure 6.**
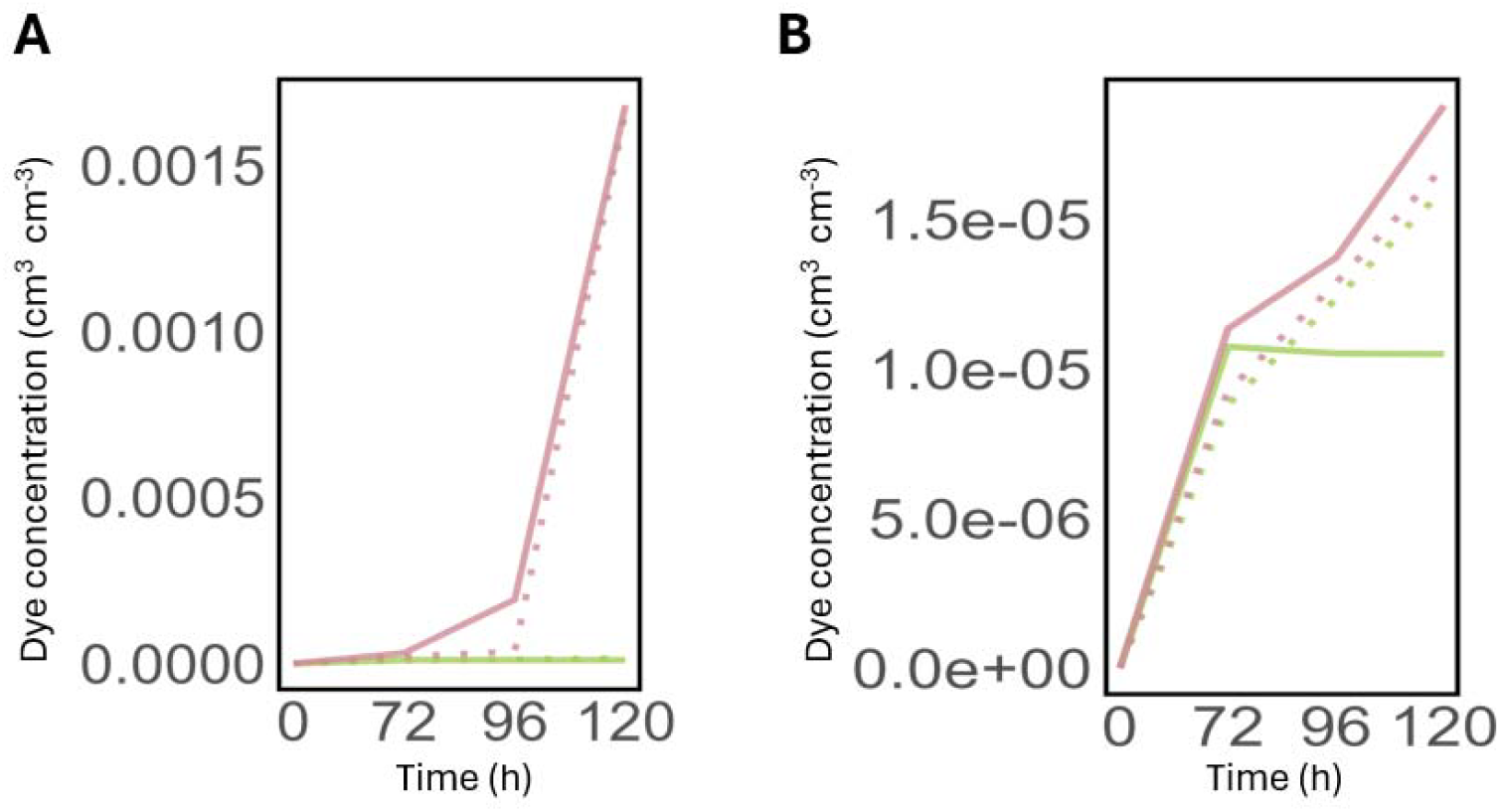
Model predictions against experimental data for dye concentration in the bottom transparent soil layer. (A) Comparison between model and experimental data for experiments when root exudates are introduced in the top layer in the tracer dye solution. EN is shown in pink and NN is shown in green. (B) Comparison between model and experimental data for experiments when root exudates are introduced dry in the middle layer. NE is shown in pink and NN is shown in green. In all plots, dotted lines represent model predictions data whereas plain lines indicate experimental observations.

## Discussion

### Water infiltration and loss of water from the rhizosphere

Plant roots absorb water stored in the soil’s pore spaces, which is subsequently replenished through irrigation, rainfall, or capillary rise. However, a substantial portion of the water input is lost before it can be absorbed by the plant. Losses are due to evaporation, deep percolation, runoff, and lateral flow and can be particularly high before canopy closure (Chen et al. 2020). In a cropping system, 20% of the evapotranspiration is typically due to bare soil evaporation, but it can exceed 50% of the total influx of water. For example, a review of the seasonal losses of water by evaporation in rain fed wheat in Australia showed that between 27% and 68% of the evapotranspiration is due to evaporation (Unkovich et al. 2018). The percentage of evaporation was between 15% and 40% in Maize in a range of water and N regimes (Hernández et al. 2015). In sandy soils, deep percolation can become the dominant source of water loss. Nassah et al. (2018) reported nearly 50% loss of water inputs lost due to water leaving the rooted zone through deep percolation.

The root system architecture, the spatial and topological arrangement of roots into a soil, is therefore a critical trait to determine the ability of a crop to extract water from the soil. First because it determines the regions of the soil where water can be extracted. When a root system is shallow, the plant can more readily acquire immobile nutrients that are strongly absorbed to soil particles (Lynch and Brown 2002). When the root system is deep, it can reach the water table and extract groundwater during drought (Lynch 2013). However, water balance in soil is very dynamic and another important role of root systems may be to intercept water, that is to distribute water into soil following a rainfall, to minimize losses from the rhizosphere. Although roots are known to facilitate preferential water flow through the soil, this function of root systems has been largely overlooked as a valuable trait to improve water management in cropping systems. Recently, mathematical models that combined water uptake with preferential flow induced by plant roots showed that root system architectures may have very different abilities to intercept rainfall water, and that root system ideotypes may vary as a function of soil type (Mair et al. 2023).

The experiments described in this study aimed to investigate the impact of root traits on water infiltration in dry soil. We used a dye tracing approach rather than infiltrometers to miniaturise and shorten the duration of the microcosm experiments. Observation of the dye also gave insights into the mechanisms of infiltration. The use of an artificial soil improved the accuracy of measurements since all the dye below the dry layer was considered in the measurement of the fluxes. The artificial soils also made experiments more manageable. Nafion^TM^, the material used as substrate, is mildly hydrophobic when dry (water contact angle of 105º (Goswami et al. 2008)), and resulted in low infiltration rates but with measurable effects after a few days of experiments. Agricultural soils which have low organic matter content usually exhibit higher infiltration rates (Basset et al. 2023), and organic matter rich soils such as peat are too hydrophobic when dry (water contact angle of up to 120º (Valat et al. 1991)).

### Physical processes involved in preferential flow induced by plant roots

We observed that root exudation was a primary factor influencing the infiltration of water through dry soil. This result suggests that surfactants found in root exudates may play a crucial role in affecting water movement within the soil. Not only root exudates are known to contain surfactants (Read et al. 2003), but measurements of water retention in rhizosphere and bulk soil have also indicated that rhizosphere solutions may exhibit lower surface tension (Whalley et al. 2005). Surfactants are also used for the irrigation of plants growing in hydrophobic soils and arid environments (Baratella and Trinchera 2018; Ogunmokun et al. 2020), which confirms that the exudation of surfactants may be a beneficial trait in certain soils.

Experiments using syringe needles indicated that changes in porosity induced by root growth may not affect significantly the infiltration of water. In a coarse and loosely packed, sand, similar to our artificial soil, root growth was shown to induce dilation of <4% and increase the porosity of about 10% over a layer of <2 mm around the root (Anselmucci et al. 2021). The Kozeny–Carman equation *K* ∝ porosity ^3^/ (1-porosity ^2^) predicts that such changes can result in an increase of 50% of the hydraulic conductivity (Schulz et al. 2019). But a 1 mm thick layer represents about 10% of the cross section of the soil. Also, dilation of the rhizosphere soil is associated with reduction of porosity further away from the root and collectively, this would likely limit the changes in the hydraulic conductivity to less than 5%. In the Young-Laplace equation, pore size and surface tension exert equal effects on capillary entry pressure (Hallett 2008). Given that root exudates have been shown to reduce surface tension by 27%, reducing it from 72 mN m^-1^ to 50 mN m^-1^ (Read et al. 2003), it is plausible for root exudation to have a stronger effect than dilation in some conditions.

### Root exudation as a mechanism to influence water availability in soil

During drought conditions, evidence suggests that both the quantity and composition of root exudates undergo significant changes (Karlowsky et al. 2018; Williams and de Vries 2020). However, it remains uncertain whether these changes reflect a plant’s response to modify the physical properties of the soil. It was suggested that for a large part, changes in root exudation result from shift in plants metabolic activity, with root exudates reflecting the most abundant root metabolites (Gargallo-Garriga et al. 2018). The lifespan of root exudates in soils is likely to be highly variable. Many of the compounds deposited in the soil by plant roots are consumed by soil microbes (Sasse et al., 2018), and it is challenging to determine the extent to which changes in soil physical properties are directly attributable to rhizodeposition, as opposed to being secondary effects arising from microbial activity. For example, soil bacteria are extraordinary producers of biosurfactants (Karanth et al. 1999; Domingues et al. 2020). They are also able to synthesize extracellular polymeric substances (EPS) which enhance soil water retention (Zheng et al. 2018; Benard et al. 2023). The reduction of surface tension typically necessitates small amounts of a single surfactant molecule, such as Lecithin found in soybeans (Read et al. 2003). Consequently, even subtle changes in the composition of root exudates could significantly impact soil properties.

It is also far from clear what the desired physical properties of the rhizosphere solution should be. While surfactant production may be desirable in low soil water content when water is trapped in small pores (Whalley et al. 2005) or in soils rich in organic matter (strong hysteresis between wetting and drying), the broader benefits are less clear. The level of organic matter in agricultural soils is minimal (Loveland and Webb 2003; Rawls et al. 2004) and heterogeneously distributed (Lehmann et al. 2008). Also, losses due to evaporation and deep percolation are extremely dependent on rainfall or irrigation patterns. Answering such questions will be challenging and is likely to require further experimental characterisation, together with the integration of mathematical modelling and simulation studies.

## Acknowledgements

We acknowledge the funding from the Spanish Ministry of Science, Innovation and Universities under the grant agreements No. PID2020-112950RR-I00 and No. PID2023-149435OR-I00 (projects MICROCROWD and BIOFLOW). This work was also supported by the European Union’s Horizon Europe under the grant agreement No.101060124 (Project Root2Res).

